# On the dynamic contact angle of capillary-driven microflows in open channels

**DOI:** 10.1101/2023.04.24.537941

**Authors:** Jodie C. Tokihiro, Anika M. McManamen, David N. Phan, Sanitta Thongpang, Terence D. Blake, Ashleigh B. Theberge, Jean Berthier

## Abstract

The true value of the contact angle between a liquid and a solid is a thorny problem in capillary microfluidics. The Lucas-Washburn-Rideal (LWR) law assumes a constant contact angle during fluid penetration. However, recent experimental studies have shown lower liquid velocities than predicted by the LWR equation, which are attributed to a velocity-dependent dynamic contact angle that is larger than its static value. Inspection of fluid penetration in closed channels has confirmed that a dynamic angle is needed in the LWR equation.

In this work, the dynamic contact angle in an open channel configuration is investigated using experimental data obtained with a range of liquids, aqueous and organic, and a PMMA substrate. We demonstrate that a dynamic contact angle must be used to explain the early stages of fluid penetration, i.e., at the start of the viscous regime, when flow velocities are sufficiently high. Moreover, the open channel configuration, with its free surface, enhances the effect of the dynamic contact angle, making its inclusion even more important. We found that for the liquids in our study, the molecular-kinetic theory (MKT) is the most accurate in predicting the effect of the dynamic contact angle on liquid penetration in open channels.

## 1. Introduction

When a fluid flows in contact with a wall, its contact angle generally differs from its static value ^1–3^. At the present time there are several approaches to predict the value of the dynamic contact angle (DCA) in the so-called viscous regime defined by Lucas, Washburn, and Rideal (LWR)^4–6^: the hydrodynamic model (HD), the molecular kinetic theory (MKT) and empirical or semi-empirical correlations based on the capillary number. The LWR law is widely used in capillary flows such as flows in channels and for flows in porous media^7,8^. This approach utilizes a static contact angle which assumes a constant contact angle throughout fluid flow. However, recent studies have shown that the use of the LWR law is applicable on a velocity-dependent basis, where in reality, the contact angle is larger than the static angle at high fluid velocities, resulting in the need for a DCA correction. This conclusion stems from the direct observation of the contact angle of spreading liquids on solid surfaces^8–11^. Dynamic contact angles are a real phenomenon. The question is how important is their effect on open channel microfluidics.

The basis for understanding the DCA has been established by, among others, de Gennes^9^ who demonstrated that, for a partial wetting, the unbalanced interfacial tension forces, *F* = γ (*cos*θ_0_ – *cos*θ_d_)— where *γ* is the liquid/air interfacial tension (unit mN/m), θ*_d_* and θ*_0_* are the dynamic and equilibrium contact angles respectively—must be compensated either by the viscous dissipation at a mesoscopic scale or by dissipation in the vicinity of the contact line or in the precursor film.

The hydrodynamic (HD) approach corresponds to the case of viscous dissipation at the mesoscopic scale resulting in so-called viscous bending near the wall (Figure 1A)^10–13^.

**Figure 1.**
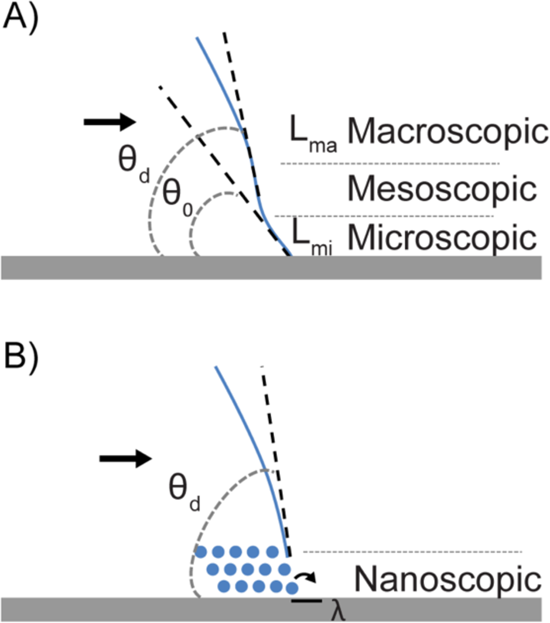
Sketches of the two main models for DCA: A, the hydrodynamic model (HD) assuming a viscous bending at the wall; B, the molecular dynamic theory (MKT) with the rolling motion of molecules at the wall.

In the HD approach, the increase in the contact angle outside of the microscopic region (L_mi_) is linked to the cubic root of the capillary number^10^

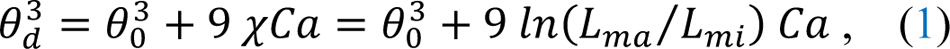

where *Ca* is the capillary number (*Ca* = μ*V*/γ, *µ* being the fluid viscosity in mPa.s and *g* the fluid surface tension; dimensionless) and *χ* is a parameter given by *χ* = *ln*(*L_ma_*/*L_mi_*), *L_ma_* being a macroscopic length scale which is of the order of the size of the meniscus and *L_mi_* is a microscopic length scale, which is the cutoff length below which the continuum theory breaks down.

On the other hand, adsorption-desorption dynamics of liquid molecules on a solid surface near the TPCL (triple phase contact line) is the basis for the molecular kinetic theory (MKT, Figure 1B). This approach was first proposed by Blake and Haynes^14^ using Eyring’s activated-rate theory of stress-modified activated rate processes^15^. According to the MKT, to an approximation, the velocity-dependent dynamic contact angle is given by^14,16,17^

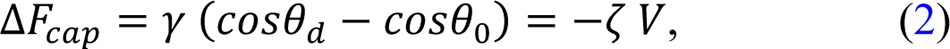

where ζ is the coefficient of contact line friction per unit length of the contact line (unit Pa.s). Strictly speaking, this simple linear form applies only near equilibrium or where the energy barriers to contact-line movement are small; but as we are interested only in the way in which a change in the dynamic contact angle may provide a correction to the viscosity-dominated flow in the channel, this form is adequate for our immediate purposes.

It was later demonstrated^16–19^ that the contact-line friction is related to the equilibrium work of adhesion, *Wa*, by

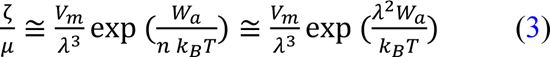

where λ is the characteristic distance of each molecular displacement (Figure 1B), *k*_B_ is the Boltzmann constant (*k*_B_ ≈ 4.14 *pN*/*nm*), *T* is the temperature (in Kelvin), *V_m_* is the molecular flow volume of the liquid, here approximated by the molecular volume. In (3), *n* is the number of solid-liquid interaction sites per unit area of the solid surface, provided these are distributed uniformly, *n* ≅ 1/λ^3^. Note that at a more fundamental level, ζ is related to the characteristic molecular frequency *K*^0^(the frequency of molecular absorption, unit MHz) and distance λ for molecular displacements in the vicinity of the triple line: ζ = *nk*_B_*T*/*K*^0^ λ ∼ *k*_B_*T*/*K*^0^ λ^3^.

Note that, in the specific case of the existence of a precursor film^20,21^ (where a thin sheet of fluid precedes the TPCL), a modified approach of MKT—called MKT-self-layering—assumes the formation of layers of molecules in the precursor film at the wall combined with the adsorption-desorption phenomenon (Figure 2)^22–24^. In this model, the wall viscosity, β, is then given by

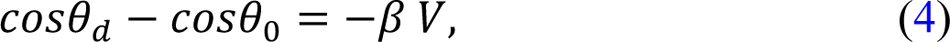

with 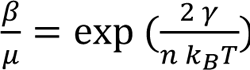, where *μ* is the viscosity of the flowing liquid, and *n* is the number of molecules per unit area in the precursor thin film, which is related to *σ* or the effective diameter of the molecules in the thin film by *n* = 1/σ^3^.

**Figure 2.**
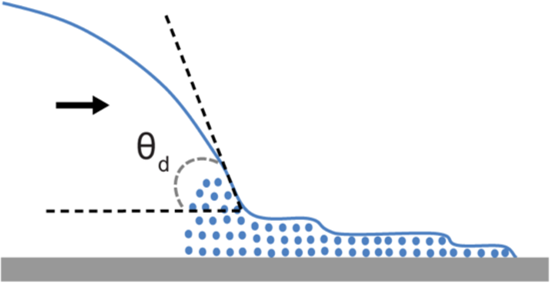
Sketch of the molecular kinetic theory – self layering approach where the precursor film is composed of layers of fluid molecules (represented as blue circles).

Finally, let us remark that empirical and semi-empirical correlations have been proposed to predict the value of the DCA. These correlations make use of the capillary number and have the form^25–30^

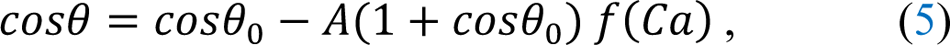

where *A* is a non-dimensional coefficient and *f* is a monotonously increasing function. In many of these correlations, the function, *f*, is of the form, *f*(*Ca*) = *ACa*^B^, where *Ca* is the capillary number, *A* ranges between 2 and 4.96, and *B* is a non-dimensional constant (ranging between 0.42 and 0.7). The term 1 + *cos*θ_0_ represents the adhesion energy, *Wa* = γ (1 + *cos*θ_0_).

In this work, we focus on the dynamic contact angle in capillary flows in open channels^31–33^. Contrary to traditional closed channel microfluidics, open channels are characterized by one or more air-liquid interfaces through the removal of at least one channel wall. Open channels have recently gained attention in the microfluidics community due to advantages such as accessibility, easy fabrication or surface treatment of the microfluidic channels, and the reduction in bubble formation during fluid addition and flow. These devices are simple to operate, necessitating only a micropipette, fluid of choice, and the open channel device. Open microfluidics has become widely used in a variety of research fields such as cell culture, protein and metabolite assays, organ-on-a-chip models, and even space applications^34–37^. To some extent, the physics of the microflow in open channels resembles that in closed channels, but some considerations must be adopted for open-channel flow. The LWR law must be modified by using an equivalent contact angle^32,33^ —the so-called generalized Cassie angle–in place of the contact angle, and an average friction length in place of the tube radius.

It is shown here that, in the early stages of the viscous regime, a dynamic contact angle (DCA) should be used and, based on our experiments, that the most accurate approach is the molecular kinetic model. Our study employs rectangular open-channels milled in poly(methyl methacrylate) (PMMA)—slightly rounded at the bottom inner corners to avoid capillary filaments^38^, and several different liquids including water, nonanol, pentanol, chloroform, FC-40, and an aqueous solution of 50% (v/v) isopropyl alcohol.

## 2. Materials and methods

### a. Channel Design and Fabrication

Four different open rectangular channels milled in PMMA have been used: channel #1 (w = 1 mm, h = 2 mm), channel #2 (w = 2 mm, h = 2 mm), channel #3 (w = 0.4 mm, h = 0.6 mm), and channel #4 (w = 0.4 mm, h = 0.4 mm). An engineering drawing of channel #1 is shown in Figure 3. Detailed engineering drawings of channels #1-4 can be found in Figure S1.1.1. Calibration markers separated by a known distance apart were milled into the device for scaling purposes during image analysis (Figure 3B). A profilometer photo of the rounded corners is shown in Figure 3B.

**Figure 3.**
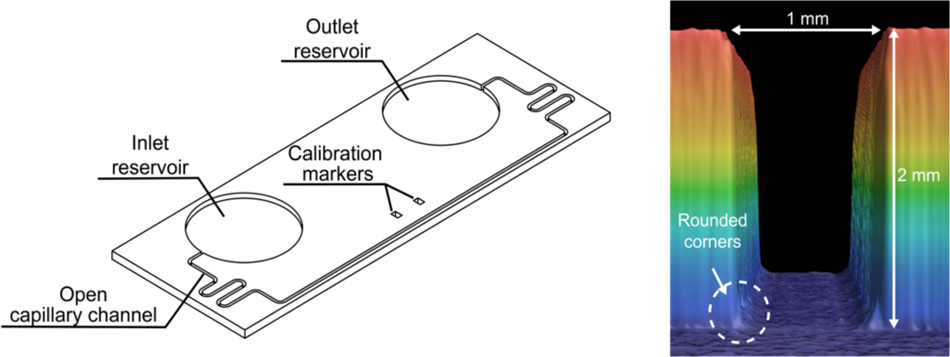
**A:** Isometric view of channel #1 milled in PMMA and **B:** profilometer cross-section of the channel with the rounded corners.

The characteristics of the channels are listed in Table 1.

**Table 1.** Characteristics of the channels.

| Channel | Width | Height | Max channel length | Wetted perimeter | Free perimeter | Friction length |
| --- | --- | --- | --- | --- | --- | --- |
| *Symbol* | $w$ | $h$ | $l$ | $p_w$ | $P_F$ | $\bar{\lambda}$ |
| *units* | $[mm]$ | $[mm]$ | $[mm]$ | $[mm]$ | $[mm]$ | $[mm]$ |
| Channel #1 | 1.0 | 2.0 | 303 | 5.0 | 1.0 | 0.22 |
| Channel #2 | 2.0 | 2.0 | 120 | 6.0 | 2.0 | 0.44 |
| Channel #3 | 0.40 | 0.60 | 110 | 1.6 | 0.4 | 0.09 |
| Channel #4 | 0.40 | 0.40 | 110 | 1.2 | 0.4 | 0.07 |

The average friction lengths (rightmost column in Table 1) representing the wall friction are first approximated by the semi-empirical formulations^33,39^ for a rounded-bottom rectangular open channel

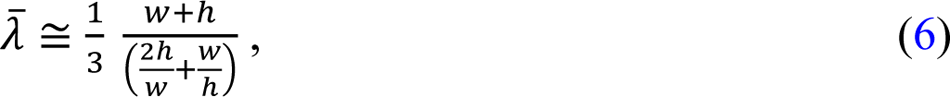

Note that for a straight-bottom rectangular open channel, an expression of the friction has been found^40–43^.

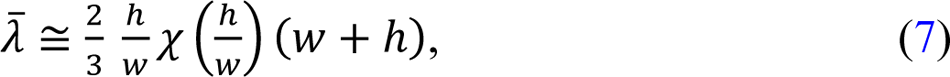

where 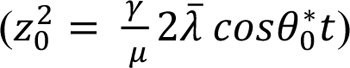.

These two formulas produce approximate values of the friction length for moderate aspect ratio rectangular open channels. Note that the second formula corresponds to the case of rectangular channels with sharp inner corners where filaments are present^44^. Hence the two formulas somewhat differ when considering the same aspect ratio channel (see SI.2).

The value of the friction length—approximated by (6) —is adjusted on the experimental plots for the flow velocity, using velocities far from the channel entrance, where the contact angle most resembles the static contact angle—as will be shown later in the text (Figure 4).

**Figure 4.**
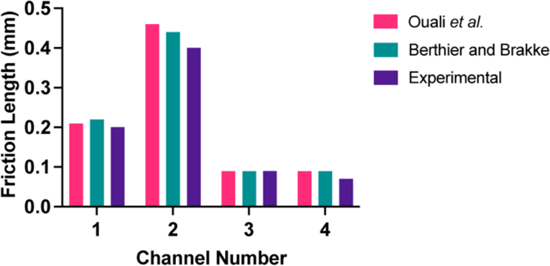
Comparison of the different approaches for the friction length: Ouali *et al.*^40^ (pink color); Berthier and Brakke^33^ (teal color); experiments (purple color).

The channels were designed using a computer-aided design (CAD) software (Solidworks 2017, Waltham, MA) and the design files were converted to a .simpl file using a computer-aided manufacturing (CAM) software (Fusion 360, Autodesk, San Rafael, CA). Channels were milled in PMMA sheets (3.175 mm thick, #8560K239; McMaster-Carr, Sante Fe Springs, CA).

To create round bottom channels, endmills with a cutter diameter of 1/32” (TR-2-0312-BN) or 1/64” (TR-2-0150-BN) were used (Performance Micro Tool, Janesville, WI). The devices were fabricated via micro-milling on a Datron Neo computer numerical control (CNC) mill (Datron, Germany). After fabrication, the channel dimensions were confirmed using a Keyence wide-area 3D measurement VR-5000 (Keyence Corporation of America, Itasca, IL). The channel bottom is estimated to have a few microns of roughness—due to the milling process— which is one magnitude below the roughness values observed by Lade *et al.* to produce substantial fluctuations in velocity^42^. Note that the effect of small roughness on open-capillary flow dynamics is not yet completely understood, since small reliefs on the wall surface slightly increase the capillary force —due to the Wenzel effect^45^ —but also slightly increase the friction force^46^. In our case, the adjustment of the friction length indirectly takes this effect into account.

### b. Solvent Prep and Physical Properties

Various solvents have been used in this study. Aqueous solvents included: deionized and distilled water (American Society for Testing and Materials Type II, HARLECO, Sigma-Aldrich, St. Louis, MO) as well as isopropyl alcohol at a concentration of 50% (v/v) in deionized and distilled water. These solvents were colored with 0.60 % yellow or 1.2 % blue food coloring. (McCormick). Organic compounds such as nonanol (Sigma-Aldrich, St. Louis, MO), pentanol (Sigma-Aldrich, St. Louis, MO), and chloroform (Fisher Scientific, Hampton, NH) have been colored with either Solvent Yellow 7 or with Solvent Green 3 (Sigma-Aldrich, St. Louis, MO) at concentrations of 0.50 mg/mL and 1.43 mg/mL respectively. FC-40 (Sigma-Aldrich, St. Louis, MO) was not colored, but tracking its travel in the channel was still feasible. In the case of water, the channel was treated by oxygen plasma using a Diener Zepto PC EX Type PB plasma treater (Diener Electronic, Germany) no more than 30 minutes prior to experimentation, to avoid contact angle relaxation^47,48^.

The physical data for these liquids are listed in Table 2.

**Table 2.** Properties of the liquids.

| | $\gamma$<br>[mN/m] | $\mu$<br>[mPa.s] | $\theta$ | w<br>[mm] | h<br>[mm] | $\cos \theta^*$ |
| --- | --- | --- | --- | --- | --- | --- |
| Water | 72 | 1.0 | 46 | 0.4 | 0.6 | 0.30 |
| Chloroform | 28 | 0.6 | 15 | 1 | 2 | 0.64 |
| 50% (v/v) Isopropyl alcohol | 30 | 3.0 | 18 | 0.4 | 0.4 | 0.46 |
| 50% (v/v) Isopropyl alcohol | 30 | 3.0 | 18 | 1 | 2 | 0.63 |
| 50% (v/v) Isopropyl alcohol | 30 | 3.0 | 18 | 2 | 2 | 0.46 |
| FC-40 | 16 | 2.5 | 17 | 1 | 2 | 0.63 |
| Pentanol | 25 | 3.7 | 13 | 2 | 2 | 0.48 |
| Nonanol | 27 | 11.5 | 13 | 2 | 2 | 0.48 |

\*Note: These values are from Kim *et al.* 2016<sup>49</sup> and Mohammad *et al.* 2014<sup>50</sup>

The values of the surface tensions and viscosities of the liquids have been taken from physical tables and the literature^49,50^. Surface tension measurements are reported as the surface tension between the air-liquid interface. The contact angles with PMMA have been measured in Lee *et al*. 2019^51^ using a Kruss DSA-25E drop shape analyzer (Kruss GmbH, Hamburg, Germany).

Contact angles are reported as the average of 5 2-µL droplet measurements collected with the Kruss ADVANCE software. The generalized Cassie angle *8\** was calculated using the static contact angle and the weighted average of the contact angle of each fluid with the wall and the atmosphere. This is defined by:

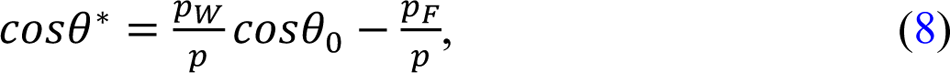

where θ*_0_* is the Young (static) contact angle, *p*_-_and *p*_=_ are the wetted and free perimeters in a cross section, and *p* = *p*_W_ + *p*_F_. In the present case, setting the cross section to a rectangle of width, *w,* and height, *h:*

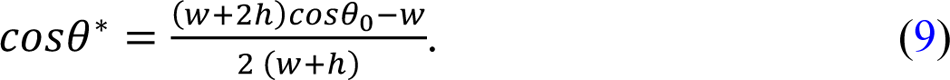

### c. Open-Channel Flow Experiments

To obtain fluid velocities, the prepared fluids were flowed through the uniform cross section channels (#1-4). In channel #1, 2.2 mL of the dyed chloroform, water, and the colorless FC-40 were added to the inlet reservoirs of individual devices. For the devices using chloroform and water, which required an extended travel distance, a refill of 300 µL of the flowing fluid was added to the inlet reservoir to minimize the effect of pressure on the fluid dynamics. A refill was not added for FC-40, since data collection stopped after the fluid front reached the first calibration marker. Data were collected for the chloroform and water experiments until the fluid front of each respective device reached the outlet reservoir. For channel #2, 2.2 mL of the dyed 50% (v/v) isopropyl alcohol, nonanol, and pentanol were added to the inlet reservoir of individual devices connected to the cross section corresponding to w = 2 mm and h = 2 mm.

Data were collected until the fluid surpassed the first calibration maker. For channel #3, 140 µL of the dyed water was added to the plasma treated device and data were collected until the fluid front reached the outlet reservoir. For channel #4, 170 µL of 50% isopropyl alcohol was added to the inlet reservoir and data were collected until the fluid front reached the outlet reservoir. For each fluid in each channel dimension, experiments were completed in triplicate (n = 3).

Videos of the progression of the flow of the solvent in the device were recorded using a Nikon-D5300 ultra-high resolution single lens reflective (SLR) camera at 60 fps. Devices were set atop a white background with a lab jack underneath. An image of the camera and device set-up can be found in Figure SI.1.2. A video frame was analyzed every 10 frames using an execution file written in Python. The distance that the fluid front had traveled was measured using ImageJ. The scale for each trial was set using the “Set Scale” function and the calibration markers on the device. The fluid front was tracked using the segmented line function and the total travel distance for each frame was measured using the “Measure” function. Data were exported as a .csv file and imported into Microsoft Excel. Calculations for fluid velocity and comparisons to the theoretical model were also conducted in Microsoft Excel.

## 3. Theoretical approach

Using the geometry of the open channel, the LWR law yields the relation between velocity, *V,* and travel distance, *z,*

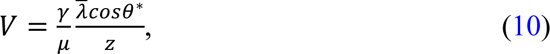

where λ is the average friction length which describes the wall friction (unit mm), θ^∗^ is the generalized Cassie angle which accounts for the effect of the free surface, *γ* is the surface tension of the liquid, and *μ* is the viscosity. Relation (10) can be rewritten in terms of the capillary number (*Ca*):

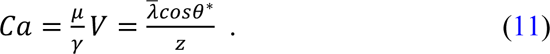

Here, the DCA depends on the value of the velocity, represented by the capillary number. Relation (11) shows that if the travel distance is sufficient and/or the channel cross section is sufficiently small (^λ^/*Z* ≪ 1), the use of the static contact angle is justified. Conversely, if this condition is not met, a DCA is present and changes the dynamics of the flow.

In this section, it is shown that the MKT approach can be applied analytically, leading to an easy modification of the extended LW law to take into account the dynamic contact angle. This observation was also made by Wu *et al.*^24^ for capillary rise, using the LWR law with an equivalent radius.

In the viscous regime, the capillary force (*F_cap_*) balances the wall friction force (whether the channel is closed or open), and

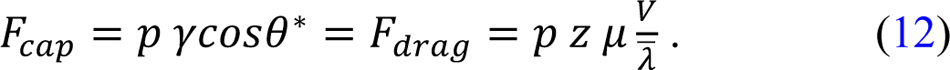

In relation (12), *z* is the marching—or travel—distance (unit mm), *p* is the total perimeter of a cross section (including the liquid/air boundary if any) (unit mm), λ is the average friction length^32^, and θ^∗^ is the generalized Cassie angle^52^. Substituting *V* = *dZ*/*dt*, relation (12) can be written as

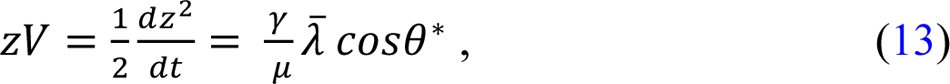

or

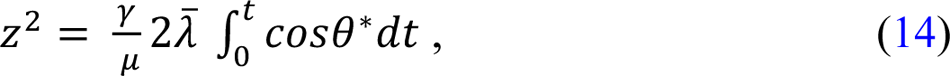

In the case where θ^∗^ is constant, the usual extended Lucas-Washburn law is retrieved. Using the expression of the generalized Cassie angle (8), relation (14) becomes

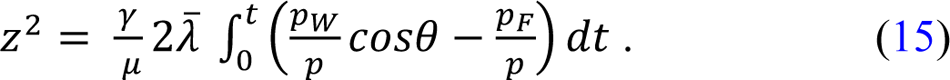

Now, let us use the MKT correlation (4) to express the dynamic contact angle

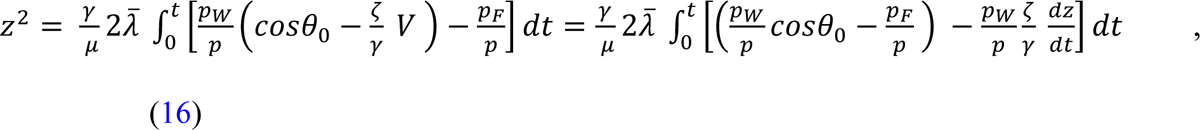

or

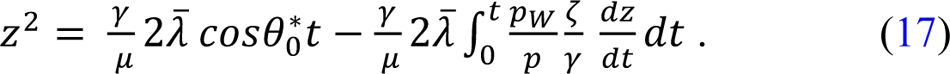

The first term in (17) is the travel distance using the static contact angle 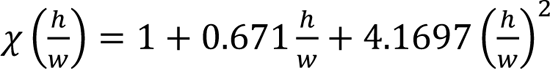, and the second term is a correction due to the dynamic contact angle. Relation (17) can be simplified as

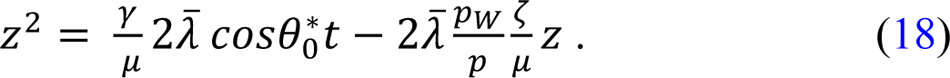

The real travel distance—accounting for the dynamic contact angle correction—is then solved using the quadratic equation:

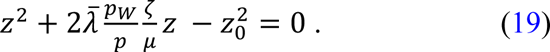

In (19), we must keep in mind that *z* and *z_0_* are functions of the time, *t*. Therefore, we finally obtain:

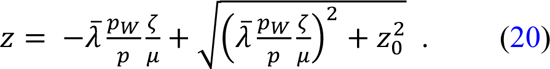

It is verified that at *t* = 0 (we neglect the evanescent inertial motion^53^), *z_0_* = 0 and *z* = 0. For a long channel, *z_0_* becomes large and *z_0_*.

Flow velocity can be easily deduced from (20) by time differentiation:

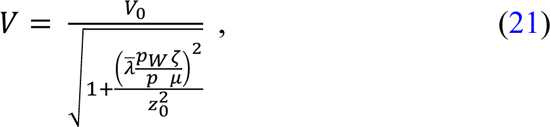

where *V_0_* is the velocity obtained using the static contact angle, *V*_0_ = *dZ*_0_/*dt*. When the travel distance increases, *z_0_* increases and the real velocity *V* converges towards the velocity, *V_0_*. Note that when *z_0_* goes to zero, z goes to zero, and relation (21) indicates that *V* goes to 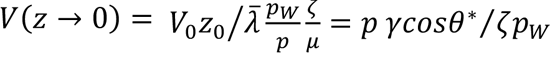, indicating that ζ acts like a viscosity associated with the triple line. However, in reality, relation (21) holds when the inertial regime ends i.e., when *Z*_0_ > *z_inertial_*

## 4. Results and Discussion

### Experimental results and comparison with model

The importance of the DCA is directly linked to the velocity of the flow. As the flow velocity in the channel is proportional to the ratio of the surface tension and viscosity V ≈ (γ/μ)/*Z*, we have categorized the fluids as “fast” and “slow” as a function of their intrinsic velocity, *V_i_* = *ψ/m*. The “fast” fluids correspond to high values of *V_i_ (V*_0_ ≥ 30 *m*/*s*) and “slow” fluids by low values of *V_i_ (V*_i_ ≤ 8 *m*/*s*).

Water and chloroform are in the first category. Figure 5 shows the dynamics of these two fluids in channels of different cross sections. Clearly, the LWR law which uses the static (Young) contact angle does not account for the flow velocity. A DCA is required to fit the experimental results as seen in Figure 5 where the experimental travel distances and velocities are closely fitted to the values predicted by the DCA theory (blue line). This DCA is obtained using ζ= 0.32 Pa.s for water and 0.20 Pa.s for chloroform.

**Figure 5.**
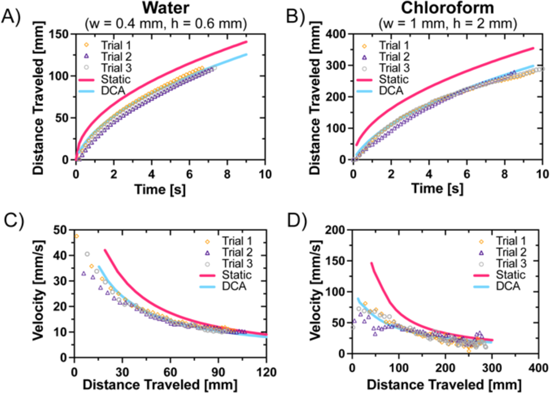
Comparison of models using the static and dynamic angles. Plots of travel distance vs. time for water in channel #3 (w = 0.4 mm, h = 0.6 mm) (**A**) and chloroform in channel #1 (w = 1 mm; h = 2 mm) (**B**): comparison between the static contact angle (pink line), dynamic contact angle model (blue line) and experiments (circles, triangles, and diamonds). Plots of velocity vs. travel distance for water (**C**) and chloroform (**D**): comparison between the static contact angle model (pink line), dynamic contact angle model (blue line), and experiments (circles, triangles, and diamonds).

The solution of 50% (v/v) isopropyl alcohol in water is located in the “intermediate” fluids, categorized by a moderate *V_i_* = *ψ/m* value. Figure 6 shows the dynamics of the flow in relatively large channels (w = 1 mm, h = 2 mm) (Figure 6A and C) compared to small cross sections (w = 0.4 mm, h = 0.4 mm) (Figure 6B and D).

**Figure 6.**
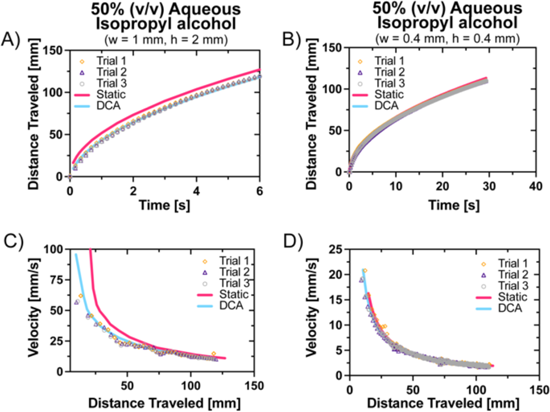
Comparison of models using the static and dynamic angles. Plots for travel distance vs. time for 50% (v/v) aqueous isopropyl alcohol in channel #1 (w = 1 mm, h = 2 mm) (**A**) and in channel #4 (w = 0.4 mm; h = 0.4 mm) (**B**): comparison between the static contact angle model (pink line), dynamic contact angle model (blue line), and experiments (circles, triangles, and diamonds). Plots of velocity vs. travel distance for 50% isopropyl alcohol in channel #1 (w = 1 mm, h = 2 mm) (**C**) and in channel #4 (w = 0.4 mm; h = 0.4 mm) (**D**): comparison between the static contact angle model (pink line), dynamic contact angle model (blue line), and experiments (circles, triangles, and diamonds).

In the first case, where a large cross section was used, a DCA is needed to account for the dynamics as shown through the experimental data clustering around the theoretical DCA travel distances and velocities (Figure 6A and C). On the other hand, the static angle (pink line) is sufficient in the second case of small cross sections (Figure 6B and D). The explanation is related to the value of the velocities, which are high in the first case, and small in the second case.

In the last case of “slow” fluids, the preceding observation is still valid. Figure 7 shows a comparison of the dynamics of pentanol and nonanol in large channels (w = 2 mm, h = 2 mm) for which the wall friction is minimal. A DCA is needed in the case of pentanol (*V_i_*∼7 m/s) (Figure 7A and C), but it is not needed for the case for nonanol (*V_i_*∼2.5 m/s) (Figure 7B and D).

**Figure 7.**
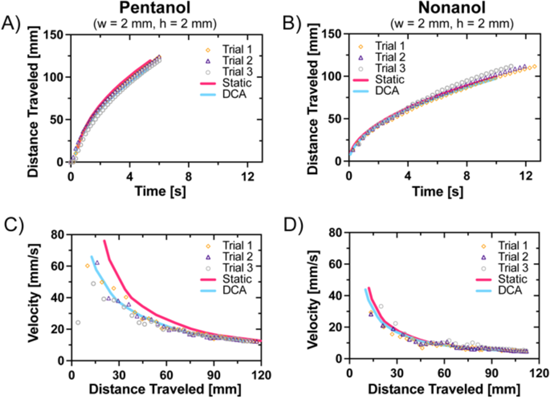
Comparison of models using the static and dynamic angles. Plots of travel distance vs. time for pentanol (**A**) and nonanol (**B**) in channel #2 (w = 2 mm, h = 2 mm): comparison between the static contact angle model (orange line), dynamic contact angle model (purple line), and experiments (circles, triangles, and diamonds). Plots of velocity vs. distance for pentanol (**C**) and nonanol (**D**): comparison between the static contact angle model (pink line), dynamic contact angle model (blue line), and experiments (circles, triangles, and diamonds).

### Comparison with other correlations

In this section, we briefly present other correlations for DCA and compare them to the MKT approach used in section 3.

Hydrodynamic correlation proposed by Hoffman and Tanner^25,26^ yields

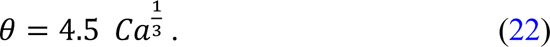

Bracke *et al.* correlation^28,29^ links the cosine of the dynamic contact angle to the capillary number:

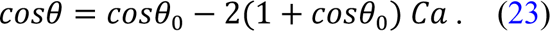

This correlation is very similar to the Seebergh and Berg^30^ correlation, while the Jiang *et al.* correlation^27^ yields:

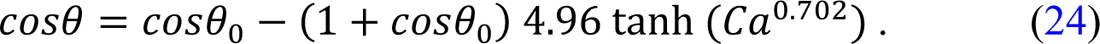

In order to investigate the effectiveness of these empirical correlations to account for the influence of a dynamic contact angle in the LWR equation, a numerical scheme was set up in a Matlab file^54^ for each correlation. A comparison of the different correlations is shown in Figure 8A for water flowing in the channel #3 (w = 0.4 mm, h = 0.6 mm). Comparison for chloroform and 50% (v/v) isopropyl alcohol are shown in SI.1.

**Figure 8.**
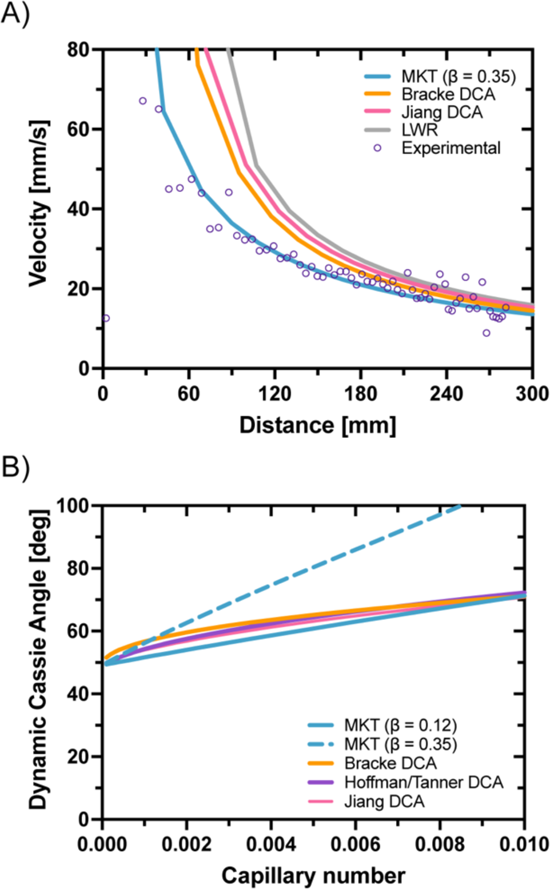
**A:** Comparison of the velocity of a water flow in channel #3 between experiments (open circles), and different correlations. **B:** Comparison between the dynamic contact angles (Cassie generalized contact angles) obtained using the different correlations described in the text in function of the capillary number. The two blue lines correspond to MKT correlations for ζ= 0.12 (blue solid line) and 0.35 Pa.s (blue dotted line).

The evolution of the DCA with the distance is plotted in Figure 8A for the capillary flow of water in channels of cross section, w = 0.4 mm and h = 0.6 mm. The different approaches— constant static contact angle, HD correlations, and MKT—are compared. The MKT approach is the only one that fits the experimental data.

The generalized DCA given by its cosine,

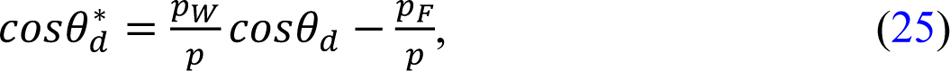

depends on the capillary number and is shown in Figure 8B (the value at *Ca* = 0 corresponds to the static Young angle) for the different correlations (Bracke *et al.*^28^, Jiang *et al.*^27^, Hoffman^25^) using the data for water. The results are very similar except for the MKT correlation with a coefficient ζ = 0.35 Pa.s.

In Figure 9A and B, the relations between the DCA and the velocity and between the DCA and the travel distance, respectively, are shown for the different liquids used in this work. The importance of taking into account a DCA is evident for the first 50 mm of the open channels.

**Figure 9.**
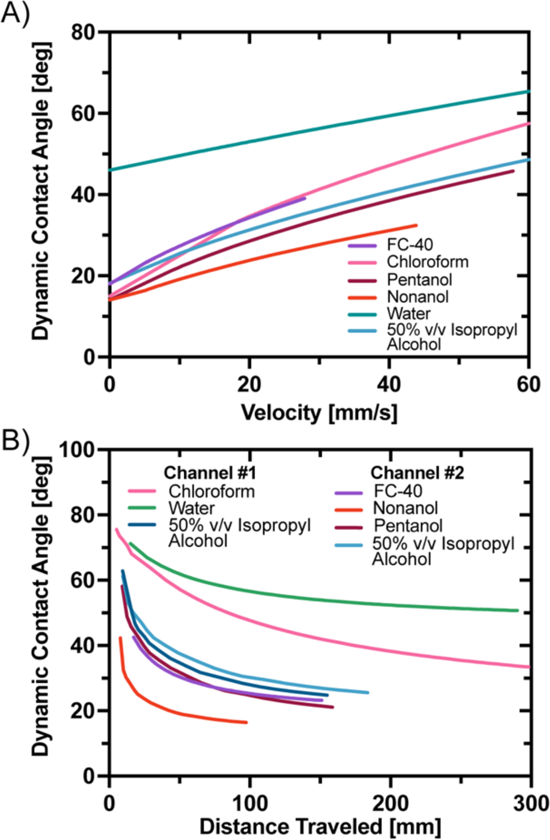
**A:** DCA vs. velocity for the 6 liquids (in the case of water, the PMMA has been treated with O_2_ plasma) and **B:** evolution of the DCA with the distance in the channel for in different cases investigated in this work.

Let us first remark that the advantage of using an open-channel configuration to assess the MKT approach is the increased sensitivity of the flow velocity to the DCA. A correction, ε, to the cosine of the contact angle has a stronger influence on the capillary force in the case of an open flow:

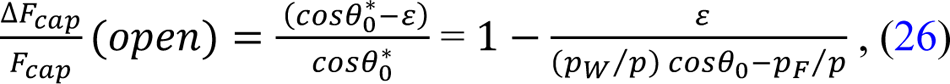

compared to

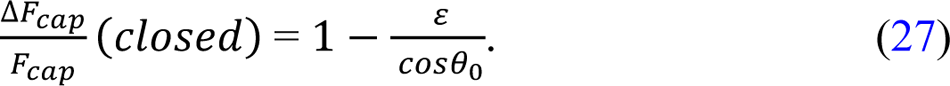

The sensitivity of an open microflow in a rectangular channel has recently been pointed out by Chang *et al.*^53^.

### Comparison between inertial and viscous regimes

A second remark is focused on the flow regimes (inertial or viscous). In the inertial regime, the velocity field is not established^55^; even if there is the effect due to the triple line friction on the contact angle, the different models presented here cannot be used. We verify that the experimental results used in this work correspond to the viscous regime. In Figure SI.2.1, the transition times and distances for the liquids and channels used in this study are calculated and listed. It is checked that the corrections associated with ζ shown in Figures 5, 6, and 7 correspond to the viscous regime and not the inertial regime. In Table 3, the inertial-viscous transitions for the different cases of Figures 5, 6, and 7 are listed. These times (t_in_) and travel distance (z_in_) transitions are located at the very beginning of the velocity vs. distance plots and do not interfere with the MKT approach (and other approaches).

**Table 3.** Transition data between inertial and viscous regimes.

| | Channel | Liquid | $t_{in}$<br>[ms] | $z_{in}$<br>[mm] |
| --- | --- | --- | --- | --- |
| Figure 5 |  |  |  |  |
|  | #3 | Water | 2.7 | 1.9 |
|  | #1 | Chloroform | 46 | 12.3 |
| Figure 6 |  |  |  |  |
|  | #1 | 50% (v/v) isopropyl alcohol | 5.7 | 2.0 |
|  | #4 | 50% (v/v) isopropyl alcohol | 0.7 | 0.4 |
| Figure 7 |  |  |  |  |
|  | #2 | Pentanol | 4.0 | 2.9 |
|  | #2 | Nonanol | 17 | 4.0 |

### Determination of molecular displacement distance

Finally, we determine the values of the molecular displacement distance (λ) of the MKT approach. Using the software, Matlab^54^, we iteratively determine the values of λ by solving the discretized equation (3) using the experimental fit of ζ

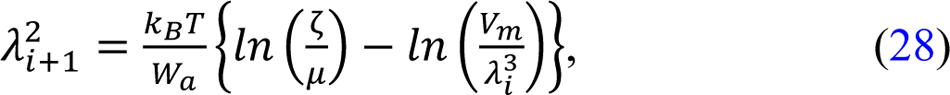

where *i* is the iteration index. Convergence is rapidly obtained in less than 10 iterations. The displacement distances are listed in the Table 4. The values of the molecular displacement frequency *K*^0^ can then be extracted using the relationship *K*^0^= *nk*_2_*T*/ζ λ. The resulting values λ and *K*^0^ are listed in Table 4. The values are usually obtained by direct measurement of the dynamic contact angle and applying the full, non-linear version of the MKT^16^. Such measurements are inherently problematic in microfluidic channels.

**Table 4.** Values of the adhesion energy, W_a_, coefficient of line friction z, molecular displacement distances λ for the different liquids and cubic root of molecular diameter *V*_m_^1/3^.

| | $W_a$ | $z$ | $\lambda$ | $V_m^{1/3}$ | $\kappa^0$ |
| --- | --- | --- | --- | --- | --- |
|  | [mN/m] | [Pa.s] | [nm] | [nm] | [MHz] |
| Pentanol | 54.6 | 0.12 | 0.45 | 0.56 | 378 |
| Nonanol | 57.1 | 0.18 | 0.30 | 0.66 | 851 |
| 50% (v/v) Isopropyl Alcohol | 62.4 | 0.12 | 0.56 | 0.39 | 196 |
| FC-40 | 31.3 | 0.22 | 0.61 | 0.84 | 82 |
| Chloroform | 53.1 | 0.20 | 0.52 | 0.51 | 147 |
| Water | 134.4 | 0.32 | 0.51 | 0.31 | 101 |

Note that, as expected, the displacement distances are in the range of 4 to 7 Angstroms, which is of the order of the cubic root of the molecular volume ( $V_m$ ): $\lambda^3 \sim V_m$ .

Note that, as expected, the displacement distances are in the range of 4 to 7 Angstroms, which is of the order of the cubic root of the molecular volume (*V_m_*): λ^3^∼*V*_m_.

Figure 10 shows that the values of the contact-line frictions ζ are in agreement with literature (see Figure SI.3.1)

**Figure 10.**
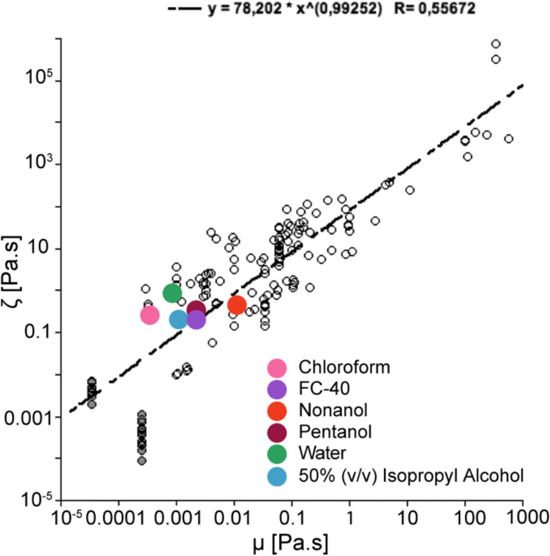
Plot of *z* vs. *μ*: Comparison of the values collated from the literature (black symbols) and the present study (colored circles)^17^. The literature references are listed in SI.3. Adapted with permission from Duvivier, D.; Blake, T. D.; De Coninck, J. Toward a Predictive Theory of Wetting Dynamics. *Langmuir* **2013**, *29* (32), 10132–10140. Copyright 2023 American Chemical Society.

## 5. Conclusion

Open channel fluidics is a rapidly expanding area of interest for both research and a wide range of potential applications. This is primarily due to advantages such as accessibility, ease of fabrication and surface treatment. However, our understanding of the flow behavior of liquids in such devices remains incomplete, particularly as regards the significance of the dynamic contact angle.

Here, we have developed a model for the influence of the dynamic contact angle on open-channel flow in the viscous regime based on the molecular-kinetic theory of dynamic wetting (MKT). This model introduces the concept of triple-line friction due to localized dissipation processes and explains the increase in the contact angle^16^. In our new work, a closed-form solution is obtained, which shows that if account is taken of the dynamic contact angle, the real travel distance is equal to that given by the generalized Lucas-Washburn-Rideal (LWR) law for open-channels^32,33^ minus a correction that correlates the dynamic contact angle with the velocity. These predictions have been validated in experiments carried out for a range of liquids in rectangular open channels made of PMMA. Overall, our experiments point to the same conclusion as Li *et al.* for ionic liquids on fluoropolymer surfaces^56^, that the molecular kinetic model accounts more precisely for the dynamic contact angle in capillary flows than the hydrodynamic model.

Significantly, the real velocity of the flow is smaller than the LWR velocity, as reported in the literature for capillary-driven microflows. This is especially important in the first few centimeters of the channel, where the velocity of flow in the early viscous regime is relatively high, and will need to be taken into account in future design and modelling of open-channel fluidic devices where the free surface enhances the effect of the dynamic contact angle.

One additional aspect of the new work is that by combining the measured coefficients of triple-line frictions ζ with a semi-empirical correlation that links the friction to the work of adhesion between the liquid and the solid^18^, we have been able to extract reasonable values for the underlying MKT parameters, the molecular jump frequency *K*^0^ and distance λ that determine ζ, without resorting to direct measurement of the dynamic contact angle, which is inherently difficult in microfluidic channels.

## Supporting information

IPA 50_Channel #4

## ASSOCIATED CONTENT

### Supporting Information

The Supporting Information and Materials are available free of charge:

- Supporting information Section SI.1.1 contains engineering drawings of the microfluidic channels and an image of the experimental observation set-up; Section SI.1.2 contains the comparisons between the other correlations; Section SI.2 provides details about the transition between inertial and viscous regimes in open channels; Section SI.3 provides a comparison with literature data for ²A. (.pdf)
- Design file: Channel #1 (.stl)
- Design file: Channel #2 (.stl)
- Design file: Channel #3 (.stl)
- Design file: Channel #4 (.stl)
- MATLAB file: Code used for correlation comparison of pentanol (.m)
- MATLAB file: Code used for correlation comparison of water (.m)
- MATLAB file: Code used for correlation comparison of chloroform (.m)
- MATLAB file: Code used for correlation comparison of IPA 50 (.m)
- Video S1: Flow of fluid through microfluidic channel (.mp4)

## AUTHOR INFORMATION

### Corresponding Authors

Jean Berthier - *Department of Chemistry, University of Washington, Box 351700, Seattle, Washington 98195, United States;*, Ashleigh B. Theberge - *Department of Chemistry, University of Washington, Box 351700, Seattle, Washington 98195, United States; Department of Urology, University of Washington School of Medicine, Seattle, Washington 98105, United States;*

### Authors

Jodie C. Tokihiro – *Department of Chemistry, University of Washington, Box 351700, Seattle, Washington 98195, United States*

Anika M. McManamen – *Department of Chemistry, University of Washington, Box 351700, Seattle, Washington 98195, United States*

David N. Phan – *Department of Chemistry, University of Washington, Box 351700, Seattle, Washington 98195, United States*

Sanitta Thongpang – *Department of Chemistry, University of Washington, Box 351700, Seattle, Washington 98195, United States*

Terence D. Blake – I*ndependent consultant, Tring, HP23 5JH, United Kingdom*

## Acknowledgements

We thank Ting Wang for assistance with figure presentation. This work was funded by National Institutes of Health National Institute of General Medical Sciences grant R35GM128648 (A.B.T.), National Center For Advancing Translational Sciences grant TL1TR002318 (J.C.T.), and the University of Washington. We also acknowledge the M.J. Murdock Diagnostics Foundry for Translational Research. The content is solely the responsibility of the authors and does not necessarily represent the official views of the National Institutes of Health or other funding sources.

