## Supporting Information for "On the dynamic contact angle of capillary-driven microflows in open channels"

### Table of Contents

| Section | | Page |
|---|---|---|
| SI.1.1 | Engineering drawings of open channels, experimental observation set-up | S2 |
| SI.1.2 | Comparison between the different correlations | S3 |
| SI.2 | Transition between inertial and viscous regimes in open channels | S4 |
| SI.3 | Comparison with literature data for $\lambda$ | S6 |

### Additional supporting information included:

Design file: Channel #1 (.stl)

Design file: Channel #2 (.stl)

Design file: Channel #3 (.stl)

Design file: Channel #4 (.stl)

MATLAB file: Code used for correlation comparison of pentanol (.m)

MATLAB file: Code used for correlation comparison of water (.m)

MATLAB file: Code used for correlation comparison of chloroform (.m)

MATLAB file: Code used for correlation comparison of IPA 50 (.m)

Video S1: Flow of fluid through microfluidic channel (.mp4)

### SI.1.1 Engineering drawings of open channels and experimental observation set-up

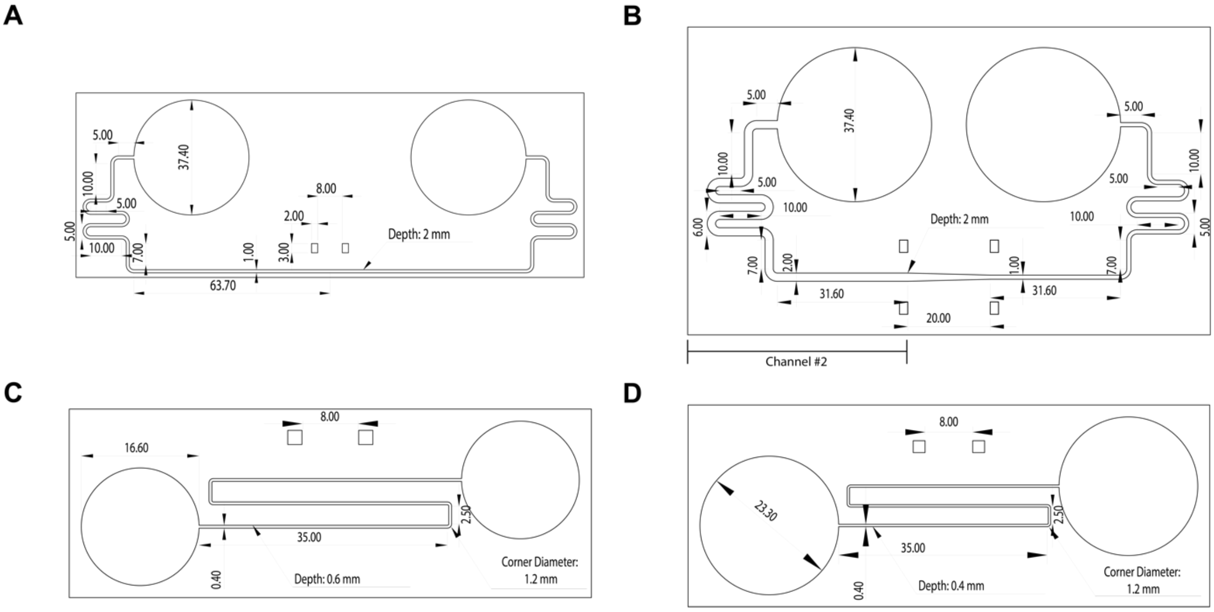


**Figure SI.1.1.** Engineering drawings for open microfluidic channels #1 (**A**), 2 (**B**), 3 (**C**), and 4 (**D**) with dimensions. Note: for the device shown in **B**, only the data resulting from the fluid flowing the left half of the device prior to the channel constriction were used in this manuscript.

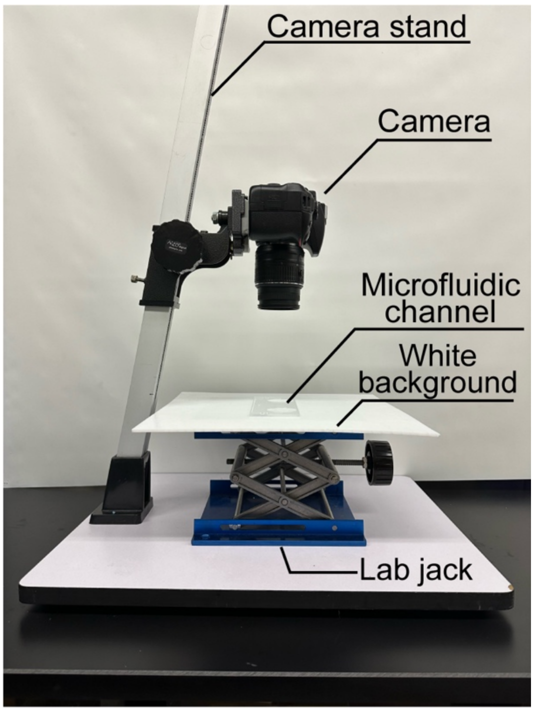


**Figure SI.1.2.** Set-up of DSLR camera and platform for experimental observation and data collection.

### SI.1.2 Comparison between different models

In this section, we compare travel distances and velocities between four models (Lucas-Washburn-Rideal (LW), correlation of *Jiang et al.,* correlation of Bracke *et al.,* MKT approach) and experimental data.

#### A. Water in an open rectangular channel (w = 0.4 mm, h = 0.6 mm)

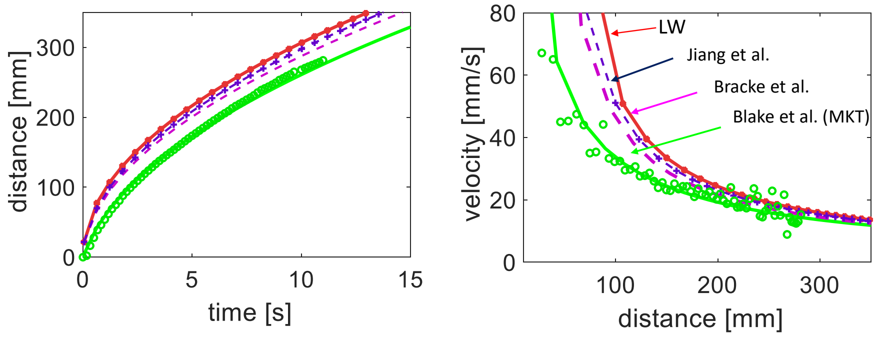


**Figure SI.1.2.1. Left:** travel distance vs. time; **right:** velocity vs. distance. Case of water. The dots correspond to the experiments. Note: The Lucas-Washburn-Rideal law is denoted as LW.

#### B. Chloroform in an open rectangular channel (w = 1 mm, h = 2 mm)

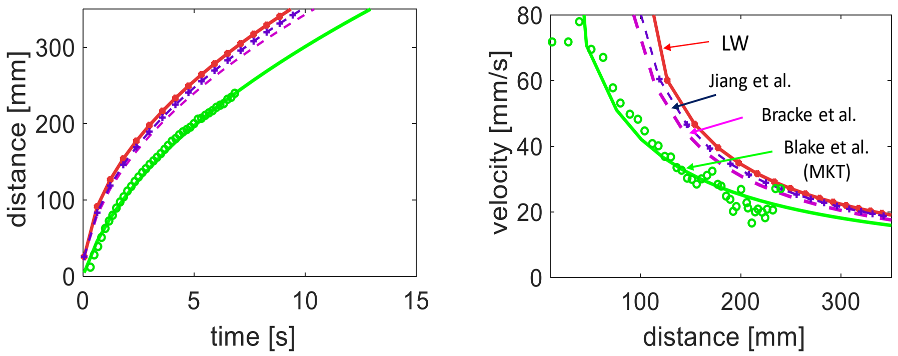


**Figure SI.1.2.2. Left:** travel distance vs. time; **right:** velocity vs. distance for the case of chloroform. Note: The Lucas-Washburn-Rideal law is denoted as LW.

#### C. 50% (v/v) isopropyl alcohol in an open rectangular channel (w = 1 mm, h = 2 mm)

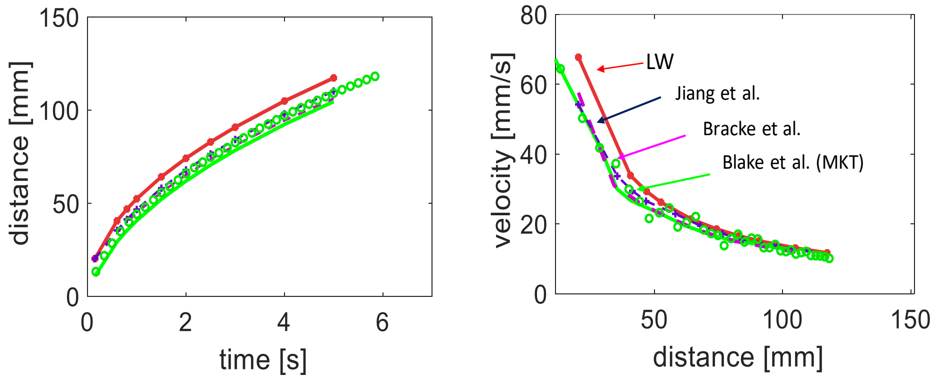


**Figure SI.1.2.3 Left:** travel distance vs. time; **right:** velocity vs. distance in the case of 50% (v/v) isopropyl alcohol. Note: The Lucas-Washburn-Rideal law is denoted as LW.

### SI.2 Transition between inertial and viscous regimes in open channels

Let us first recall the Bosanquet equation for the capillary motion. In the case of an open capillary flow, the Bosanquet equation makes use of the generalized Cassie angle $\theta*$:

$$\rho\, S_C \frac{d\,(z\,\dot{z})}{dt} + \frac{\mu\, p\, z\, \dot{z}}{\bar{\lambda}} - p\,\gamma\, cos\theta^* = 0. \qquad \text{(SI.2.1)}$$

The cross-sectional area, $S_c$, is delimited by the wetted walls and the flat free surface (zero pressure). The term $\bar{\lambda}$ is average friction length. Let us note $a = \mu\, p/\bar{\lambda}\, \rho S_C$ (unit 1/s) and $b = p\gamma cos\theta^*/\rho S_C$ (unit m$^2$/s$^2$). The division of all the terms of (S1.2.1) by $\rho\, S_C$ yields:

$$\frac{d}{dt}\left(z\frac{dz}{dt}\right) = b - a\left(z\frac{dz}{dt}\right). \qquad \text{(SI.2.2)}$$

Assuming $z$ = 0 at $t$ = 0, time integration leads to the differential equation in $z^2$:

$$\frac{dz^2}{dt} + az^2 = 2bt. \qquad \text{(SI.2.3)}$$

A second integration yields the solution,

$$z^2 = \frac{2b}{a}\left[t - \frac{1}{a}(1 - e^{-at})\right]. \qquad \text{(SI.2.4)}$$

The inertial regime occurs when the wall friction can be neglected at the beginning of the motion, so that (SI.2.3) becomes

$$\frac{dz^2}{dt} = 2bt. \qquad \text{(SI.2.5)}$$

Time integration of (SI.2.5) yields

$$z = \sqrt{\frac{p\,\gamma\, cos\theta^*}{\rho\, S_C}}\; t = \sqrt{b}\, t. \qquad \text{(SI.2.6)}$$

The inertial velocity $V_{in}$ is then constant, leading to

$$V_{in} = \sqrt{\frac{p\,\gamma\, cos\theta^*}{\rho\, S_C}} = \sqrt{b}\,. \qquad \text{(SI.2.7)}$$

The end of inertial regime occurs when the term $(1 - e^{-at})$ in (SI.4) becomes negligible, i.e. $t \sim\ 1/4a$. Then, the inertial-viscous transition time is

$$\tau_{inertial} \sim \frac{1}{4}\,\frac{\bar{\lambda}\,\rho\, S_C}{\mu\, p} = \frac{\bar{\lambda}\,\rho\, D_H}{16\,\mu}, \qquad \text{(SI.2.8)}$$

where $D_H$ is the hydraulic diameter defined by $D_H = 4\,\frac{S_C}{p}$. The travel distance at the end of inertial regime is then

$$z_{inertial} \simeq \sqrt{b}\,\tau_{inertial}\,. \qquad \text{(SI.2.9)}$$

The inertial distance depends both on friction length and Cassie angle. The different values for the inertial times and inertial distances are listed in Table SI.2.1.

| *Liquids/channels* | #1 $t_{in}$ [ms] | #1 $z_{in}$ [mm] | #2 $t_{in}$ [ms] | #2 $z_{in}$ [mm] | #3 $t_{in}$ [ms] | #3 $z_{in}$ [mm] | #4 $t_{in}$ [ms] | #4 $z_{in}$ [mm] |
|---|---|---|---|---|---|---|---|---|
| Pentanol | 3.4 | 0.79 | 4.2 | 1.48 | 0.48 | 0.17 | 0.37 | 0.13 |
| Nonanol | 1.17 | 0.29 | 3.13 | 0.54 | 0.16 | 0.06 | 0.12 | 0.05 |
| 50% (v/v) isopropyl alcohol | 0.49 | 0.35 | 1.33 | 0.66 | 0.70 | 0.08 | 0.52 | 0.06 |
| FC-40 | 10.6 | 1.37 | 29.1 | 2.60 | 1.53 | 0.31 | 1.16 | 0.23 |
| Chloroform | 38.3 | 6.97 | 103.0 | 13.1 | 5.42 | 1.55 | 4.14 | 1.18 |
| Dodecane | 9.65 | 2.35 | 26.0 | 4.41 | 1.36 | 0.52 | 1.04 | 0.39 |
| Toluene | 22.7 | 5.42 | 61.2 | 10.19 | 3.21 | 1.20 | 2.45 | 0.91 |
| Water | 15.4 | 4.36 | 41.7 | 7.62 | 2.18 | 0.95 | 1.67 | 0.68 |

Table SI.2.1. Inertial times [ms] and distances [mm]

In Figure SI.2.1, we have indicated the values of the inertial distance in three open rectangular channels, and eight different liquids (including the six liquids that we use in this work). We remark that the dissociation of the Bosanquet equation for motion (SI.2.1) results in overestimated cutoff times. In reality, a progressive transition occurs between the two regimes.

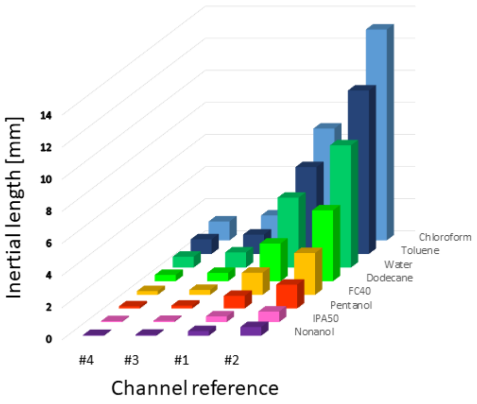


**Figure SI.2.1.** Different inertial distances for the four rectangular open-channels and eight liquids.

### SI.3. Comparison with literature data for λ

The molecule volumes corresponding to the different molecules used in the study are obtained from tables of molar mass and listed in Table SI.3.1.

| | Molar mass [g/mol] | Density [g/mL] @ 25°C | Molecular volume [nm$^3$] |
|---|---|---|---|
| Nonanol | 144.3 | 0.830 | 0.29 |
| Pentanol | 88.2 | 0.815 | 0.18 |
| 50% (v/v) isopropyl alcohol | 40.0 | 1.100 | 0.06 |
| FC-40 | 650.0 | 1.850 | 0.59 |
| Chloroform | 119.0 | 1.483 | 0.13 |
| Water | 18.0 | 1.0 | 0.03 |

**Table SI.3.1.** Molecular volume of the different liquids

Inserting these molecular volumes, $V_m$, and the values of the coefficient, $\zeta$, fitted on the experimental results in the MKT relation,

$$\frac{\zeta}{\mu} = \frac{V_m}{\lambda^3} \exp\left(\frac{W_a}{n\,k_B T}\right) \cong \frac{V_m}{\lambda^3} \exp\left(\frac{\lambda^2 W_a}{k_B T}\right), \quad \text{(SI.3.1)}$$

and we obtain the implicit relation linking $\lambda$, $\mu$, and $W_a$:

$$ln\left(\frac{V_m}{\lambda^3}\right)\frac{\lambda^2 W_a}{k_B T} \cong ln\left(\frac{\zeta}{\mu}\right). \quad \text{(SI.3.2)}$$

The solution of (SI.3.2) is achieved by a numerical scheme. Equation (SI.3.2) can be written as:

$$\lambda^2 = \frac{k_B T}{W_a}\left\{ln\left(\frac{\zeta}{\mu}\right) - ln\left(\frac{V_m}{\lambda^3}\right)\right\} = \frac{k_B T}{W_a}\left\{ln\left(\frac{\zeta}{\mu}\right) - lnV_m + 3ln\lambda\right\}. \quad \text{(SI.3.3)}$$

Then, the iterative explicit approach is:

$$\lambda_{i+1}^2 = \frac{k_B T}{W_a}\left\{ln\left(\frac{\zeta}{\mu}\right) - ln\left(\frac{V_m}{\lambda^3}\right)\right\} = \frac{k_B T}{W_a}\left\{ln\left(\frac{\zeta}{\mu}\right) - lnV_m + 3ln\lambda_i\right\}, \quad \text{(SI.3.4)}$$

where *i* is the iteration index and produces the solution, $\lambda$. Convergence is rapidly obtained in less than 10 iterations. We obtain the values of $\lambda$ listed in Table SI.3.2.

| | Molecular volume [nm$^3$] | $W_a$ [mN/m] | $\lambda$ [nm] |
|---|---|---|---|
| Nonanol | 0.29 | 57.1 | 0.30 |
| Pentanol | 0.18 | 54.6 | 0.45 |
| IPA50 | 0.06 | 62.4 | 0.56 |
| FC40 | 0.59 | 31.3 | 0.61 |
| Chloroform | 0.13 | 53.1 | 0.52 |
| Water #1 | 0.03 | 116.6 | 0.49 |
| Water #2 | 0.03 | 113.7 | 0.50 |
| Water #3 | 0.03 | 113.7 | 0.50 |
| Water #4 | 0.03 | 110.7 | 0.51 |
| Water #5 | 0.03 | 101.0 | 0.54 |

**Table SI.3.2.** Work of adhesion and average distances of molecule displacement

We can compare these values of $\lambda$ with the data found in the literature and collated by Duvivier, Blake and De Coninck in reference [SI.3.1]. In Figure SI.3.1, we observe that the values of the average distance of each displacement $\lambda$ are located precisely in the cloud of data collated in the literature.

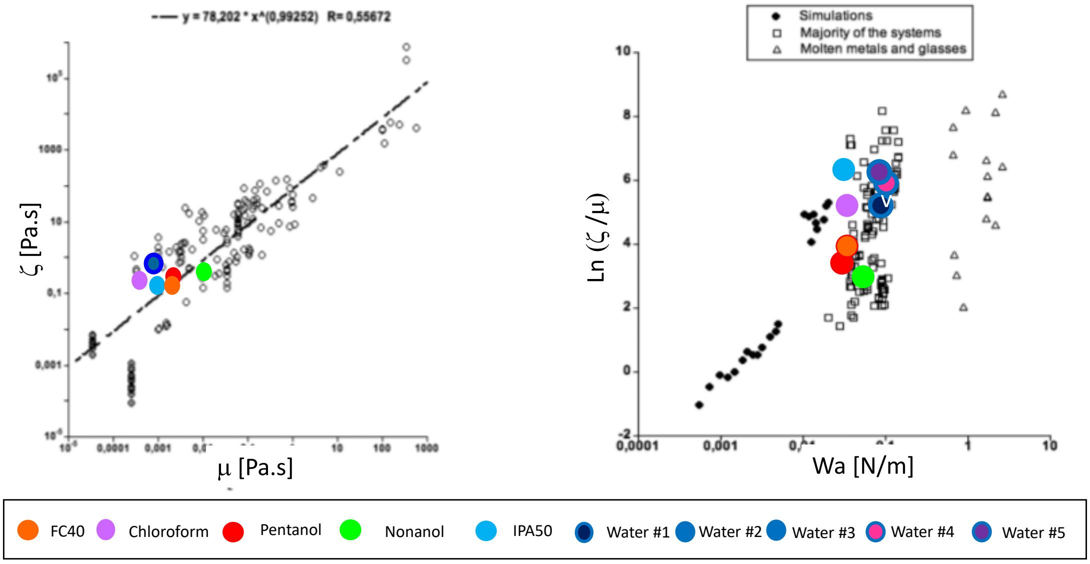


**Figure SI.3.1. Left:** coefficient of line friction vs. fluid viscosity; **right:** ln(ζ/η) vs. work of adhesion (Wa). The black symbols (circles, diamonds, triangles) correspond to literature data extracted from [SI.3.1]. Adapted with permission from D. Duvivier, T.D. Blake, J. De Coninck 2013 Toward a predictive theory of wetting dynamics, *Langmuir* **29**(32),10132-10140. Copyright 2023 American Chemical Society. Note: 50% (v/v) aqueous isopropyl alcohol is denoted as IPA50.
